# Interpersonal brain synchronization with instructor compensates for learner’s sleep deprivation in interactive learning

**DOI:** 10.1101/2020.04.03.022954

**Authors:** Yafeng Pan, Camille Guyon, Guillermo Borragán, Yi Hu, Philippe Peigneux

## Abstract

Recent advances shifted the focus on single-brain functioning toward two-brain communication during learning interactions, following the demonstration that interpersonal brain synchronization (IBS) can track instructor-learner information exchange. Here, we investigated (*i*) whether sleep deprivation (SD) that potentially impacts both social interactions and learning abilities modulates IBS, and (*ii*) conversely whether and to what extent IBS might compensate for SD-related learning deficits. Instructors (always with regular sleep, RS) were asked to teach numerical reasoning strategies to learners (either SD or RS), during which the activity of both brains was simultaneously recorded using functional near-infrared spectroscopy (fNIRS). SD learners initially performed below their baseline level, worse than RS learners, but learning improvement was comparable between RS and SD conditions after learning with the instructor. IBS within the instructor-learner dyads was higher in the SD (vs. RS) condition in the left inferior frontal cortex. In addition, clustered IBS (estimated by nonnegative matrix factorization) was correlated with performance improvement. Finally, Granger Causality analyses revealed biased causality with higher instructor-to-learner than learner-to-instructor directionality in brain signal processing. Together, these results indicate that SD-related learning deficits can to some extent be compensated via interactions with an instructor, as reflected by increased IBS and preserved learning ability. It suggests an essential role of the instructor in driving synchrony between teaching and SD learning brains during interactions.

## 1. Introduction

Morning sleepiness is known to interfere with educational efficiency in teenagers and young adults, due to altered psychosocial and life-style circumstances but also to the maturation of biological processes regulating sleep/wake systems [1,2]. Insufficient sleep [3–8] and/or poor sleep quality [9,10] in children, adolescents and young adults are associated with cognitive dysfunctions, mood changes, and social deficits that may affect learning and academic performance.

According to the empathic deficit hypothesis [11], sleep deprivation (SD) affects the ability to recognize and categorize others’ emotions [12], and reduces the individual’s self-perceived emotional intelligence by affecting the ability to be empathetic towards others [13]. Moreover, SD is associated with impaired emotional reactivity [14] and control [15] – both being relevant for sharing others’ emotional state and social interaction. In this respect, empathic deficits are probably amongst the main factors that impede social exchange [3], which ultimately have the potential to impede the relation between an instructor and a learner during interactive learning [16].

Notwithstanding, SD-related social interaction deficits might to some extent be compensated by recruiting resources beyond those utilized after a normal night of sleep, a compensatory phenomenon usually accompanied by stronger or more extended brain activity [17] (but see e.g. [18] for decreased brain activity after SD) and increased intrinsic brain connectivity [19–21]. Hence, according to the compensatory recruitment hypothesis [17], the sleep-deprived brain would exhibit compensatory neural activity that enable relatively preserved behavioural performance, including learning. It is worth noticing that at least partially shared brain networks encompassing fronto-temporo-parietal regions subtend compensatory recruitment [17,20] and social cognition [22,23]. It suggests that compensatory mechanisms might support social exchanges in resilient individuals after one night of SD. Notwithstanding, whether and how compensatory single-brain findings apply/generalize to a dual-brain framework [24] remain unclear.

Besides an involvement of specific brain networks, functional near-infrared spectroscopy (fNIRS) – based hyperscanning studies have evidenced brain-to-brain coupling mechanisms underpinning social interactions [25–29]. Interpersonal brain synchronization (IBS) was identified using this hyperscanning approach, in which brain cortical dynamics can be simultaneously recorded in two interacting participants (also known as “second-person neuroscience” [24,30]). Using a naturalistic interactive learning paradigm, we previously showed that IBS is able to track learning interactions within instructor-learner dyads, and correlates with learning outcomes [26,28]. Furthermore, increased IBS was found to facilitate transfer of social information [31]. Improved information transfer may eventually promote better learning outcomes [26,32]. However, it remains unclear whether sleep deprivation known to affect both social cognition and learning abilities exerts an effect of IBS, and conversely whether socially based IBS might compensate for SD-related deficits in learners.

In this study, we recorded simultaneous brain activity using fNIRS, and computed IBS between an instructor and a learner during a numerical reasoning learning session [29] under regular-sleep (RS) and sleep-deprived (SD) conditions, counterbalanced. In the RS condition, both the instructor and learner spent one night of sleep at home before testing in the morning. In the SD condition, the learner was totally sleep deprived for one night under controlled conditions before testing in the morning, whereas the instructor was normally rested. Due to the interactive nature of face-to-face learning, we anticipated an IBS between instructor and learner. Considering the hypotheses discussed above, IBS could be decreased after SD due to deficient empathic abilities (for a discussion of the positive links between IBS and empathy, see [33,34]), alternatively enhanced if a compensatory mechanism develops. The latter compensatory hypothesis is inspired by both previous single-brain studies (e.g., [17,20,22,23]) and emerging two-/multi-brain findings showing that IBS selectively increases in individuals with behavioural/social deficits (e.g., [35,36]). Additionally, to better understand the functional significance of IBS at the neurophysiological level, we explored IBS-behaviour relationships (i.e., whether IBS is associated with learning outcomes), directionality coupling (i.e., whether it is the instructor who mostly synchronizes with the learner, or vice versa), and intrinsic connectivity in each partner.

## 2. Methods

### 2.1. Participants

Eighteen female adults (age 22.72 ± 1.99 years, range 20–28 years) completed the whole experiment. They were recruited through a public announcement at the Université Libre de Bruxelles (ULB, Belgium). We tested only female participants in order to mitigate inter-individual and inter-dyad variability, in accordance with recent hyperscanning studies [26,37]. Recent use of psychiatric or hypnotic drugs, poor sleep quality (Pittsburgh Sleep Quality Index > 8 [38]), and high caffeine consumption (> 3 cups/day) excluded subjects from participation in this study (two additional participants were screened). Participants gave written informed consent prior to this experiment approved by the ULB-Erasme Hospital Ethics Committee (Reference P2018/284). All participants received monetary compensation for their participation.

All participants but two were right-handed (Edinburgh Handedness Inventory, Oldfield, 1971), in good health with no history of sleep, neurologic, or psychiatric disorders, and exhibited below cut-off scores levels for anxiety (State-Trait Anxiety Inventory-French version [39]), depression (Beck Depression Inventory-Short Form [40]), empathy (Interpersonal Reactivity Index [41]) and usual fatigue (Brugmann Fatigue Scale [42]). They also had satisfactory usual sleep quality (Pittsburgh Sleep Quality Index [38]) and neutral or moderate chronotype (Morningness-Eveningness Questionnaire [43]) (see **Table 1**).

**Table 1.** Questionnaire scores of the study sample.

|  | Mean | Standard Deviation | Range | Cut-off scores |
| --- | --- | --- | --- | --- |
| Handedness | 65.82 | 51.81 | -75–100 | / |
| State-Trait Anxiety Inventory (STAI) |  |  |  |  |
| *STAI - Trait* | 28.74 | 3.64 | 21–34 | 40 |
| *STAI - State* | 29.10 | 1.72 | 23–40 | 40 |
| Beck Depression Inventory | 1.53 | 1.88 | 0–3 | 9 |
| Brugmann Fatigue Scale | 3.90 | 2.36 | 0–11 | 12 |
| Pittsburgh Sleep Quality Index | 3.32 | 1.56 | 0–5 | 7 |
| Morningness-Eveningness Questionnaire | 53.11 | 8.88 | 30–69 | 69 |
| Interpersonal Reactivity Index (IRI) |  |  |  |  |
| *IRI - Perspective Taking* | 19.00 | 5.81 | 7–28 | 30 |
| *IRI - Fantasy* | 18.11 | 4.90 | 9–27 | 30 |
| *IRI - Empathic Concern* | 20.28 | 4.59 | 12–28 | 30 |
| *IRI - Personal Distress* | 12.17 | 5.58 | 2–22 | 30 |

From the pool of 18 participants, two with instructor training (Educational Sciences) for at least 2 years were selected and assigned as instructors. Only two instructors were used in order to make the teaching style as similar as possible across dyads [44]. The remaining 16 participants were learners, randomly assigned to the 2 instructors. Thus, each instructor had to teach 8 learners in a face-to-face format, both in the RS and SD conditions (see next section), resulting in 16 instructor-learner dyads in total. This target number of participant dyads was determined by an a priori power analysis based on effect sizes reported in the literature from our previous study using an interactive learning paradigm [26]. Using reported effect sizes (Cohen’s *d*s > 0.79), the power analysis (using the *pwr* package in R [45]) indicated that a sample of 15 participant dyads would be sufficiently powerful (at a level of 0.80) to detect an effect of this size. Along these lines, previous hyperscanning studies using the instructor-learner interactive learning paradigm have typically used sample sizes of around 12 – 15 participant dyads [26,46,47].

### 2.2. Experimental protocol

In a repeated-measures crossover counterbalanced design, the learners (n = 16) took part in two experimental conditions (**Fig. 1A**) separated by a one-week interval, counterbalanced: (*i*) once after a night of regular sleep at home (i.e., regular-sleep, RS), and (*ii*) once after 24 h of sleep deprivation (SD).

**Fig. 1.**
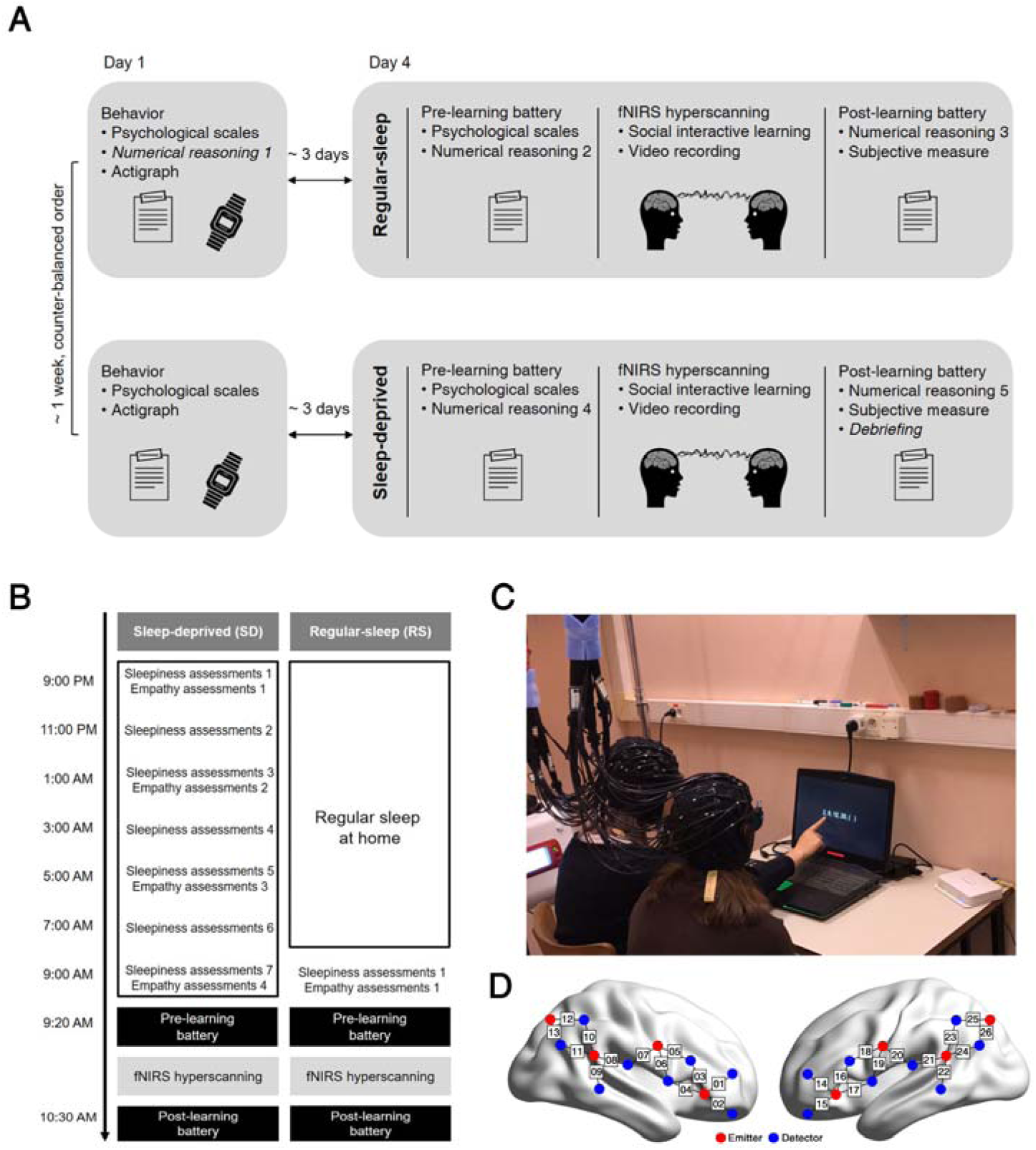
Experimental setting. (A) Schematic description of the experimental protocol. The experiment spanned on 4 days, two per condition (sleep-deprived vs. regular-sleep, SD vs. RS). Test sessions were separated by at least one week. (B) Detailed timetable in the SD and RS conditions. (C) Illustration of the fNIRS - hyperscanning experimental setup. (D) Optode emitters [red dots] and detectors [blue dots] located on bilateral fronto-temporo-parietal regions, both for the instructor and the learner.

Each experimental condition (SD or RS) included two behavioural sessions. On Day 1, participants completed a set of psychological scales (i.e., Pittsburgh Sleep Quality Index, STAI–Trait, Morningness-Eveningness Questionnaire, Beck Depression Inventory, Edinburgh Handedness Inventory, and Interpersonal Reactivity Index), and were administered a numerical reasoning test to evaluate their baseline proficiency level. At Day 4 three days later, they participated in the fNIRS-hyperscanning experimental session either after a night of regular sleep (RS) at home or total sleep deprivation (SD) in the laboratory. In this fNIRS-hyperscanning experimental session, they had first to complete self-report questionnaires (STAI – State and Brugmann Fatigue Scale) and a numerical reasoning pre-learning assessment. Then, the instructor taught numerical reasoning strategies (see below) to the learner in a one-on-one interactive format while their cortical activity was simultaneously recorded, and their interactions videotaped. Immediately after the scanning session, participants were administered a post-learning numerical reasoning assessment.

Day 1 and Day 4 for the second condition held at least one week later were identical as the first condition but for the status of the night preceding the fNIRS-hyperscanning session (SD or RS), and the fact that no baseline numerical reasoning evaluation was administrated. RS and SD sessions were counterbalanced across participants. In both RS and SD conditions, normal rest-activity patterns (7 to 9 hours of sleep) were monitored using actimetry (i.e., a wrist-worn device monitoring motor activity; ActiGraph wGT3X-BT Monitor, Pensacola, FL) and subjective sleep logs (Stanford Sleepiness Scale [48]) for 3 consecutive days before the fNIRS-hyperscanning session (**Fig. 1A**). Actigraphy data were analyzed using ActiLife 6 software [49] and the Cole Kripke algorithm [50]. Participants were also specifically instructed not to engage in daytime naps and to abstain from alcohol and caffeine during this period, including throughout the SD night.

In the RS fNIRS-hyperscanning session, participants came to the laboratory at 9:00 AM after a night of regular sleep at home. In the SD fNIRS-hyperscanning session, participants arrived at the laboratory at ∼8:50 PM the day before. Starting from 9:00 PM. and every 2 h, participants were administered sleepiness and empathy questionnaires, and a psychomotor vigilance task (PVT) (details below). The SD session enrolled two participants in a same night. They were allowed access to the Internet, books, and movies with low to moderate emotionality levels; physical activity was restricted to short walks and food intake to a small sandwich at ∼3:00 AM. Water was available ad libitum. At ∼ 9:20 AM in both conditions, participants performed the fNIRS - hyperscanning interactive learning session (with pre-learning, learning and post-learning phases, **Fig. 1A&B**).

Instructors (n = 2) were never sleep deprived and participated in the scanning/learning conditions after regular sleep at home (normal sleep patterns were monitored using actimetry). They had to complete a battery of psychological scales (i.e., Pittsburgh Sleep Quality Index, STAI-Trait, Morningness-Eveningness Questionnaire, Beck Depression Inventory, Edinburgh Handedness Inventory, and Interpersonal Reactivity Index) during the first meeting. Besides, they were asked to complete self-report questionnaires (STAI-State and Brugmann Fatigue Scale) as well as the sleepiness and empathy assessments at 9:00 AM every time when arriving at the laboratory.

### 2.3. Interactive learning task

Participants were taught numerical reasoning strategies, i.e. to find the hidden rules and relations within a digit sequence. For example, for a given digit sequence “1, 3, 5, (), 9”, the hidden rule is that all digits in the sequence are odd numbers that differ by the constant of “2”; as a result, “7” is the correct answer. Numerical reasoning items were extracted from the Chinese Civil Servants Administrative Professional Knowledge Level Tests (CCSAPKLT), a national standard guidebook. CCSAPKLT was designed to measure and improve a variety of cognitive abilities, entailing numerical reasoning, in young adults. It was previously used in an fNIRS-based hyperscanning study [29]. Learners in our study had never been exposed previously to the CCSAPKLT.

Prior to the formal experiment, the two instructors received a teaching training to ensure consistent strategies when interacting with the learners during the experimental teaching sessions. They were given 8 numerical reasoning instances (selected from CCSAPKLT) and the teaching script, and asked to prepare their teaching at home for one week. They then had to demonstrate teaching to the experimenter in a one-on-one manner, and received feedbacks until their performance was deemed satisfactory by the principal investigator (Y. P.).

During the fNIRS hyperscanning interactive learning phase, the instructor taught several numerical reasoning strategies to the learner face-to-face (**Fig. 1C**). The instructor was not informed of the learner condition (but it is likely that they realized the sleep state of the learner, as sleep-deprived learners looked exhausted). The task procedure was as follows: (*i*) the instructor presented an example on a computer screen; (*ii*) the learner read and thought about the problem for approximately 20 seconds; and then (*iii*) the instructor guided the learner to find the hidden rule according to the approach described in the script using a questions and answers (Q&A) approach. The learner’s numerical reasoning performance was evaluated at the beginning of the experiment (baseline level) and immediately before and after the fNIRS hyperscanning (for a total of five tests).

To determine the learning material, 50 four-choice items were selected from CCSAPKLT’s test bank. To create five tests with equal difficulty levels (i.e., baseline, SD-pre, SD-post, RS-pre, and RS-post), we asked 10 additional participants (not involved in the main experiment) to solve the problems, and used their scores to determine the difficulty level of the 50 items. We then rejected 10 items based on the following criterion: (*i*) confusing expressions based on pilot participants’ feedback; (*ii*) highest (>70%) and lowest levels (<30%) of accuracy (to avoid potential ceiling/flooring effects). The 40 remaining items were pseudo-randomly split into five testing sets of 8 items each. Difficulty levels did not significantly differ between the five testing sets (*t*s < 1.44, *p*s > 0.16). During baseline and pre-and post-learning tests, participants were allowed a maximum of 20 minutes (the exact time for each learner was recorded) to complete their testing set.

The numerical reasoning performance was assessed using the *Efficiency* score (adapted from [51]):

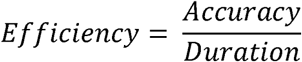

which was measured as a ratio between accuracy (defined as the ratio between the correct items and the total number of items) and duration (defined as the time in seconds necessary to complete the items). This measure was chosen to control for the potential tradeoff between accuracy and duration. Statistical analyses on numerical reasoning performance (i.e., *Efficiency*) were conducted using nonparametric Wilcoxon tests since data were not normally distributed (Shapiro-Wilk test, *p* = 0.04).

### 2.4. Sleepiness and empathy assessments

#### 2.4.1. Sleepiness assessments

##### Objective sleepiness

To assess SD-related changes in objective alertness across the night, we administered the 10-min version of Psychomotor Vigilance Task (PVT, [52]) every 2 hours from 9 PM. In the PVT, participants are instructed to press a key as fast as possible whenever a millisecond countdown appears in the middle of a computer screen. Stimuli were randomly presented with an inter-stimuli interval ranging 2 to 10 seconds. PVT response speed (i.e., reciprocal reaction time = mean 1/RT) and lapses (i.e., number of RTs > 500 ms) were the primary outcomes [53].

##### Subjective sleepiness and fatigue

To assess subjective changes in sleepiness over the SD night and before the task in SD and RS conditions, the Stanford Sleepiness Scale was administered every 2 hours from 9 PM. Participants had to choose the statement that defined them best from 7 options ranging from “feeling active and vital; wide awake” to “sleep onset soon; lost struggle to remain awake.” Participants also completed a 10 cm visual analog scale (VAS) to rate their fatigue (i.e., ‘How tired do you feel?’), ranging from not very tired (left) to very tired (right).

Sleepiness assessments (PVT, Stanford Sleepiness Scale, and fatigue VAS) were obtained every 2 h during the SD night (9:00 PM to 9:00 AM) and in the morning of the RS session (∼9:00 AM) (see **Fig. 1B**).

#### 2.3.2. Empathy assessments

##### Objective empathy

To track changes in empathy after SD using objective measurements, we administered a modified version of the Multifaceted Empathy Test (MET) [11]. This test uses 120 color pictures of people, selected from the International Affective Picture System [54]. From this set of images, we created 5 parallel versions of the task (i.e., to be administered at the 4 testing sessions across the whole night in the SD condition (9:00 PM, 1:00 AM, 5:00 AM, and 9:00 AM) and once in the RS condition (9:00 AM); administration order counterbalanced). The five subsets entailed 24 pictures each, each comprising 8 pictures for each valence: positive, negative or neutral, matched for arousal. In the task, the 24 images were presented four times, resulting in 96 trials in total. At each trial, participants had to answer a specific question that aimed at measuring either (*i*) cognitive empathy (i.e., ‘how much could you feel about the thoughts of this person?’), (*ii*) direct emotional empathy (i.e., ‘how strong is the emotion you feel about emotions of this person?’), (*iii*) indirect emotional empathy (i.e., ‘how calm/aroused does this picture make you feel?’), or (*iv*) a mere image valence judgement (i.e. ‘how would you judge this image?’ positive/negative/neutral). Each question was presented at first for 4 s, followed by a fixation cross (1 – 3 s) and then the image stimulus. When presented with the stimulus, they had to respond as fast as possible, at maximum within 10 seconds. Participants responded to cognitive and emotional empathy questions by using a reduced version of the Self-Assessment Manikin [55] valence scale which consists of four figures, ranging from calm and not concerned to anxious and very concerned. The sum of ratings of cognitive empathy and emotional empathy was calculated as an index of general empathy.

##### Subjective empathy

To track the subjective empathy changes following SD, a 10 cm VAS was used to rate participants’ empathy (i.e., ‘how much do you feel about the emotions of others?’), ranging from not very much (left) to very much (right).

Empathy assessments (MET and empathy VAS) were obtained every 4 h during the SD night (from 9:00 p.m. to 9:00 AM) and in the morning of the RS session (∼9:00 AM) (see **Fig. 1B**).

##### SD-related empathic deficit

A potential SD-related empathy deficit was calculated by subtracting objective/subjective empathy levels at ∼9:00 AM in the SD condition from that in the RS condition.

### 2.5. fNIRS data acquisition

The instructor and the learner sat side-by-side in front of a computer in a silent room (**Fig. 1C**). Brain imaging data were collected from the instructor and the learner simultaneously using a multichannel BrainSight NIRS system (V2.3b12, Rogue Research Inc., Canada). The configuration of the optodes featured 8 light emitters and 16 detectors, clustered over the fronto-temporo-parietal regions based on previous studies showing that these regions are associated with social cognition and interaction [56]. The detectors were located at a distance of approximately 3 cm from the emitters. Each pair of emitters and detector formed one channel, resulting in a total of 13 channels measured over each hemisphere (**Fig. 1D**). Probe set locations were checked and adjusted to ensure consistency within the instructor-learner pair, and across pairs. The spatial position of the optodes was set up using a 3D coordinates system coupled with a Polaris localization device. Since no individual structural magnetic resonance images (MRI) were available for our participants, MNI coordinates of fNIRS channels were determined using a probabilistic registration method [57–59]. This method utilizes MRI stored in a reference database and probabilistically registers fNIRS channel positions onto a standard brain template. The probabilistic registration consisted of four steps [59]. First, we measured positions for channels and reference points (real-world space) using the 3D digitizer. Four reference points were used: Nz (i.e., nasion), Cz (international 10/20 system), AL and AR (i.e., left and right preauricular points). Second, we applied an affine transformation of the fNIRS channel coordinates on the participant’s head (real-world space) to the reference heads in the database (MNI space). Third, we projected head surface points onto their corresponding cortical surfaces in MNI space. Finally, the cortically projected channel positions for each participant were integrated to generate the most likely coordinates in MNI space.

Absorption of near-infrared light at two wavelengths (685 and 830 nm) was measured with a sampling rate of 10 Hz. Based on the modified Beer-Lambert Law, changes in oxy-hemoglobin (HbO) and deoxy-hemoglobin (HbR) concentrations were obtained by measuring fNIRS light absorption changes after transmission through the cortical tissue. In this study, we focused on HbO concentrations only, since HbO was reported to be a sensitive indicator to reveal changes in the regional cerebral blood flow [60] and of high signal-to-noise [7,61], which has been successfully used in the field of social neuroscience to evidence IBS in recent hyperscanning studies [26,28,29,37,62].

Data collection started with a baseline 5-minute resting-state phase during which both participants were required to focus on a same fixation point on the computer screen, while keeping still and avoiding unnecessary movements. The interactive-learning phase immediately followed for an approximate duration of 13 to 18 minutes.

### 2.6. fNIRS data analyses

#### 2.6.1. Pre-processing

Data collected during rest and interactive learning (task) phases were pre-processed as follows. First, 30-second signal blocks were removed from the initial and ending rest and task phases to ensure steady state periods. Next, a Correlation-Based Signal Improvement method based on the negative correlation between HbO and HbR concentrations was applied to further reduce motion artifacts and improve signal quality [63]. Moreover, a principal component analysis was applied on continuous fNIRS data to separate the neuronal from global components [64].

#### 2.6.2. Interpersonal brain synchronization (IBS) measurement

Pre-processed data were then analyzed using wavelet transform coherence (WTC) to explore the relationship between the two fNIRS time series generated by each participant in the dyad. WTC analysis was computed using a standard MATLAB package (http://grinsted.github.io/wavelet-coherence/; see also [65] for more information). WTC function for a pair of signals *i*(*t*) and *j*(*t*) was defined as follows:

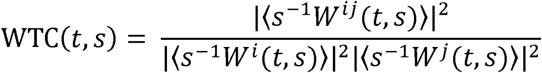

where *W* denotes a complex coefficient matrix calculated by the continuous wavelet transform with the Morlet wavelet as the mother function. This *W* matrix contains information about both instructor and learner signals’ amplitude and phase. The cross-wavelet transform *W*^*ij*^ = *W*^*i*^*W*^\**j*^ of the two signals is calculated with * indicating the complex conjugate. Moreover, *t, s*, and ⟨·⟩ represents the time, wavelet scale and smoothing operation in time and scale, respectively [65]. WTC values range from 0 (totally unsynchronized) to +1 (perfectly synchronized). WTC values were converted into Fisher-z values. IBS between instructor and learner was estimated by WTC as described in previous studies [26,29]. Considering 26 channels per participant, a 26 × 26 IBS matrix was generated for each dyad. The IBS matrices were further rendered over 3D head models using a plot function [66]. The resulting map was corrected for multiple comparisons using the false discovery rate (FDR) [67], thresholded at a 0.05 significance level.

##### Task-evoked IBS

As a first step to assess whether our interactive learning task evoked IBS, we performed the IBS analysis across all channel combinations and all conditions. To do so, IBS was averaged across time and all channel combinations in each dyad. The averaged IBS was then compared between the resting-state phase and the interactive-learning phase using paired sample *t*-tests. Comparisons were conducted for each frequency band within the 0.01 – 1 Hz range, including almost all frequencies reported in previous fNIRS hyperscanning studies [26,28,68]. The resulting *p* values were corrected using FDR. This data-driven analysis evidenced task-evoked IBS in frequencies ranging from 0.16 to 0.19 Hz (see **Fig. 3A**). That is, IBS was significantly larger in the task than the rest phase in the frequency of 0.16 – 0.19 Hz. This frequency band was thus chosen as our frequency of interest (FOI) for further analyses. It also excluded undesired effects from physiological noises [e.g., cardiac pulsation (∼1 Hz), respiration (∼0.2 – 0.3 Hz) and Mayer waves (∼0.1 Hz)].

**Fig. 2.**
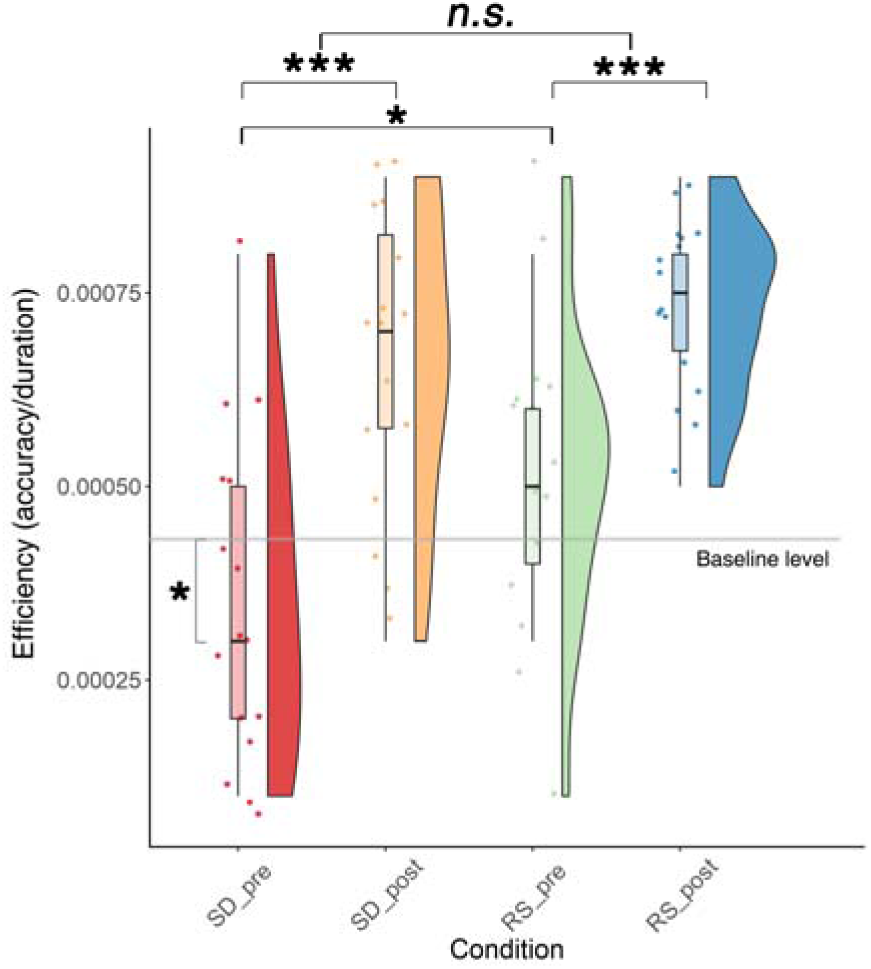
Learning performance. Numerical reasoning learning performance was quantified using *Efficiency* index (i.e., accuracy/duration). Raincloud plots [74] displaying summary data (box plot), distribution (probability density plot), and raw observations (one point = one learner). The horizontal grey line represents the baseline level (as assessed at the learners’ first visit). SD_pre: pre-learning in the sleep-deprived condition; SD_post: post-learning in the sleep-deprived condition; RS_pre: pre-learning in the regular-sleep condition; RS_post: post-learning in the regular-sleep condition. \**p* < 0.05, \*\*\**p* < 0.001.

**Fig. 3.**
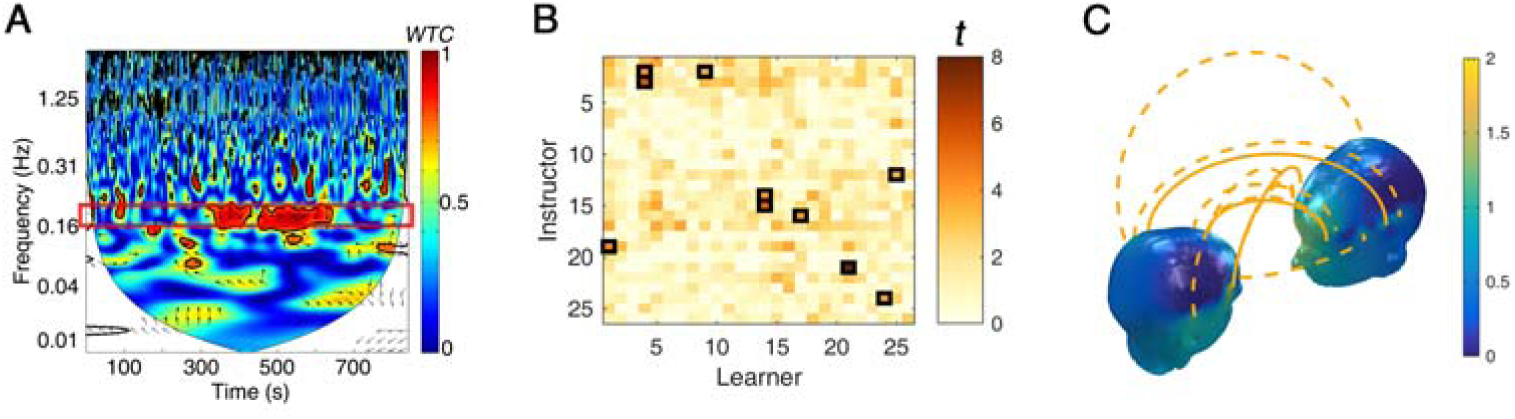
Task-evoked interpersonal brain synchronization (IBS). (A) IBS estimated by Wavelet Transform Coherence (WTC) from a representative dyad. The red border line denotes the frequency band of interest (0.16 – 0.19 Hz). (B) IBS matrix at 0.16 – 0.19 Hz. The x-axis represents channels from learners whereas the y-axis represents those from instructors. The colour indicates *t* value. Black rectangles highlight significant differences between task and rest phases. (C) Task-evoked (task vs. rest) IBS at 0.16 – 0.19 Hz across the whole sample (illustrated by orange lines; dash line indicates FDR corrected *p* < 0.05, the solid line represents FDR corrected *p* < 0.01). Head colour reflects the number of significant IBS links.

##### State-related differences in IBS

In a second step, we compared the IBS in the SD and RS conditions. We averaged the IBS within the FOI (i.e., 0.16 – 0.19 Hz.) in each condition, and computed an index of task-related IBS by subtracting IBS during rest from that during the task (i.e., *IBS*_*task-related*_ = *IBS*_*task*_ – *IBS*_*rest*_). Two complementary analyses were then conducted on task-related IBS. First, we aimed at determining condition-specific IBS [i.e., contrasting IBS values with the null value hypothesis (IBS = 0) for each channel combination using one-sample *t*-tests separately in the RS and SD conditions]. Second, we evaluated the condition-related IBS [i.e., contrasting IBS values between RS and SD conditions for each channel combination using paired-sample *t*-tests]. *P* values derived from both measures were corrected with FDR, respectively.

##### IBS validation

To confirm that the detected IBS was specific to real instructor-learner dyads, we pseudo-randomly re-paired signals from all participants into 16 new shuffled dyads (always random dyads) and re-conducted the IBS analysis. This shuffling procedure was conducted 1,000 times.

##### Clustered IBS

In the next step, IBS matrices in the SD and RS conditions were clustered using nonnegative matrix factorization (NMF) [69]. NMF is an unsupervised learning approach used to extract meaningful information from multi-dimensional data such as IBS arrays [34]. To achieve stable results, we conducted the NMF with 1,000 runs. As our primary interest, channel combinations associated with significant IBS during interactive learning in the SD condition were selected as features; as a control, we used the same features in the RS condition. The number of the cluster (i.e., factorization rank) was set to 3. This parameter was determined by considering the smallest value at which the decrease in the residual sum of squares (RSS) is lower than the decrease of the RSS obtained from reshuffled data [70], in accordance with recent recommendations [34]. The Brunet version of NMF was applied. The *coef* function implemented in the *NMF* package [69] in R 3.5.1 was used to estimate the cluster loadings.

##### IBS-behavior correlation

Pearson correlational analyses were performed to test the relationship between clustered IBS values and numerical reasoning performance (as well as empathy scores) in both SD and RS conditions. Resulting *p* values were FDR-corrected. Statistical analyses were conducted using MATLAB (version 2016b, MathWorks Inc., Natick, MA) and SPSS (version 18.0, Chicago, IL, USA) software.

#### 2.6.3. Directional coupling

We further asked about coupling directionality (i.e., whether it was mostly the instructor who synchronized with the learner or the other way around) during interactive learning after sleep loss, using a Granger causality estimation toolbox (https://www.dcs.warwick.ac.uk/~feng/causality.html). After pre-processing (see section above), which made the time series relatively stationary (as confirmed by the augmented Dickey-Fuller test) [71], time series were normalized using z-transforms (i.e., converting task phase data into z-scores using the mean and standard deviation of rest phase data). As previously reported [26], clean time series from adjacent channel combinations exhibiting significant IBS condition-related differences were averaged as a region of interest (ROI). We used the average signal from significant channel combinations, as opposed to the individual signals from each significant channel combination, in order to aggregate the effect and mitigate global systemic noises. The mean pair-wise conditional Granger Causality of the pair of time series was computed for both directions [i.e., from learner (L) to instructor (I), L → I, and from instructor to learner, I → L]. Latent variables were the averaged signals from channel combinations associated with non-significant IBS in both instructors and learners. The model order was set to 12 based on the Bayesian information criterion [72]. Ljung-Box Q-tests confirmed that there was no significant autocorrelation in the residuals. Because the data were not normally distributed (Shapiro-Wilk test, *p* = 0.03), nonparametric Wilcoxon tests were used to compare the difference between the two directions in each condition (SD and RS), Bonferroni corrected for multiple comparisons.

#### 2.6.4. Within-individual, seed-based intrinsic brain network

As a complementary analysis, we also explored seed-based intrinsic brain network to assess whether interactive learning would modify individual within-brain synchronization in the instructor and/or the learner. Seed-based intrinsic brain synchronization was estimated using the aforementioned WTC method. Inferior frontal cortex (IFC, MNI coordinates: *x* = -52, *y* = 36, *z* = -12) was selected as our seed of interest for two main reasons: (*i*) significant IBS condition-related differences were identified in the IFC in this study (see Results), and (*ii*) our previous work on instructor-learner interactions evidenced significant IBS in the IFC [26]. We calculated the synchronization between IFC (channel 15, closest to the MNI coordinates) and the remaining channels (channels 1–14 & 16–26) in each participant of dyads (25 channel combinations in every participant). The resulting synchronization matrices revealed intrinsic brain connectivity patterns that were strengthened/weakened during interactive learning. Three contrasts of interest were computed. First, task-related IBS (IBS during task minus that during rest) was contrasted against the null hypothesis (IBS = 0) to evidence interactive learning-related intra-brain synchronization; second, it was contrasted across conditions (SD vs. RS) in each participant of dyads aiming at identifying condition-related differences; third, it was contrasted across roles (instructor vs. learner) separately in the SD and RS conditions to explore potential role-related differences. The resulting *p*-values from these contrasts were controlled using FDR multiple-comparisons correction. Directional coupling and intrinsic brain networks were visualized using the BrainNet Viewer [73].

## 3. Results

### 3.1. Behavioral data and task performance

#### 3.1.1 SD-related changes in sleepiness and empathy

Actimetry and subjective sleep logs validated that participants were well rested for the 3 consecutive days before the fNIRS - hyperscanning session (asleep 8.19 ± 0.86 hours for each night, rising at ∼ 8:16 ± 0:57 AM). There were no significant differences in sleep duration and wake-up time for the three consecutive days before testing between the two sleep conditions (*p*s > 0.12). These results rule out the possibility that irregular sleep before RS and SD night might have impacted upon sleepiness and empathy.

We then assessed both objective (PVT) and subjective (SSS and VAS-sleepiness) measures of sleepiness and vigilance in learners. All scores were significantly altered in the SD vs. RS conditions when tested at 9:00 AM: PVT-response speed, mean ± standard deviation, 2.68 ± 0.44 vs. 3.42 ± 0.33; PVT-lapses, 11.69 ± 2.32 vs. 1.50 ± 0.60; SSS, 5.46 ± 1.99 vs. 2.33 ± 0.85; VAS-sleepiness, 8.83 ± 2.97 vs. 3.27 ± 2.16; *p*s < 0.002, as well across the SD night in the SD condition (9:00 AM vs. 9:00 PM); PVT-response speed, 2.68 ± 0.44 vs. 3.33 ± 0.35; PVT-lapses, 11.69 ± 2.32 vs. 1.69 ± 0.48; SSS, 5.46 ± 1.99 vs. 2.50 ± 0.90; VAS-sleepiness, 8.83 ± 2.97 vs. 3.67 ± 2.32; *p*s < 0.002. These results confirm that sleepiness and vigilance parameters were altered in learners in the SD condition.

Empathy in learners was also impaired following SD, as evidenced using both objective (MET) and subjective (VAS-empathy) assessments. VAS-empathy scores at 9:00 AM were decreased after SD vs. RS (5.50 ± 2.78 vs. 7.17 ± 1.67, *p* = 0.004) as well as over the night in the SD condition (9:00 AM vs. 9:00 PM; 5.50 ± 2.78 vs. 7.75 ± 1.83, *p* = 0.0001). The decline in MET scores was significant for the negative stimuli (9:00 AM, SD vs. RS; 8.21 ± 0.70 vs. 9.14 ± 0.51, *p* = 0.0005; 9:00 AM vs. 9:00 PM in the SD condition, 8.21 ± 0.70 vs. 9.22 ± 0.61, *p* = 0.0003), but not for neutral and positive stimuli (*p*s > 0.10).

As for instructors, who always had regular sleep during the experiment, we also collected their scores of sleepiness and empathy assessments each time when they arrived at the laboratory in the morning (9:00 AM). All scores (obtained when instructors taught RS learners) were not significantly different from those of learners in the RS condition: PVT-response speed, mean ± standard deviation, 3.06 ± 0.17 vs. 3.42 ± 0.33; PVT-lapses, 2.20 ± 2.14 vs. 1.50 ± 0.60; VAS-sleepiness, 3.31 ± 2.27 vs. 3.27 ± 2.16; VAS-empathy, 7.88 ± 1.71 vs. 7.17 ± 1.67; MET-negative, 9.35 ± 1.04 vs. 9.14 ± 0.51, *p*s > 0.05. Moreover, no significant difference was detected between scores obtained when instructors taught SD learners and scores obtained when they taught RS learners, *p*s > 0.28. These results indicate that our instructors were likely to be well rested and comparable with our RS learners across experimental sessions.

#### 3.1.2 Numerical reasoning learning performance

Summary, distribution and raw data are visualized (**Fig. 2**). Numerical reasoning learning performance across learners was quantified using *Efficiency*, which was measured as the ratio between accuracy and duration (see Methods). We performed a series of planned contrasts using nonparametric Wilcoxon tests instead of ANOVAs, since the data were not normally distributed. First, we tested whether performance at pre-learning in the SD and RS conditions differed from baseline performance levels as assessed the first day of the experiment and whether they differed from each other. In the SD condition, performance at pre-learning (median ± median absolute deviation, 0.00030 ± 0.00017) was inferior to baseline (0.00045 ± 0.00013), *p* = 0.04. In the RS condition, no significant difference was evidenced between pre-learning and baseline level (0.00050 ± 0.00014; *p* = 0.28). At pre-learning numerical reasoning performance following SD was significantly worse than that following RS, *p* = 0.04.

Second, we assessed the effect of the interactive learning session by contrasting performance at pre- vs. post-learning. Performance was superior at the post- than the pre-learning phase both in the SD (0.00070 ± 0.00016) and RS (0.00075 ± 0.00009) conditions, *p*s < 0.002. To compare learning-related improvement in performance between SD and RS conditions, we computed a differential (delta) value by subtracting pre-learning from post-learning performance. Learning improvement (delta) was not significantly different between the SD (0.00030 ± 0.0.00017) and the RS conditions (0.00025 ± 0.00014; *p* = 0.18).

These results indicate that numerical reasoning performance significant declined following SD, but similarly improved through the interaction with the instructor during the learning session in RS and SD conditions.

### 3.2. IBS during interactive learning

IBS (estimated by WTC, see Methods) was used to analyse fNIRS data. In a first-pass analysis, IBS was calculated at each channel combination across all conditions for each participant. To focus on task-related synchronized brain activity, IBS during the task (interactive learning) was computed against IBS during the rest (baseline) phase. Results are shown in **Fig. 3**. Increased IBS was evidenced in the 0.16 – 0.19 Hz frequency range (**Figs. 3A**) in a widespread cortical network encompassing inferior frontal and superior temporal areas and the temporo-parietal junctions, *t*s > 3.59, corrected *p*s < 0.05 (**Figs. 3B&C**).

### 3.3. SD-related changes in IBS

Having confirmed that interactive learning evokes distributed IBS across participant dyads, we sought to determine whether there was unchanged, impaired or compensatory IBS after SD. Analyses were conducted in SD and RS conditions separately. In the SD condition, learning (task)-related IBS significantly increased (one sample *t*-test against 0 value) in a wide fronto-temporo-parietal network, peaking at superior temporal and inferior frontal regions, *t*s > 4.57, corrected *p*s < 0.04 (**Fig. 4**, left column). In the RS condition, learning (task)-related IBS was found in the superior temporal cortex only, *t*_15_ = 6.23, corrected *p* = 0.01 (**Fig. 4**, middle column).

**Fig. 4.**
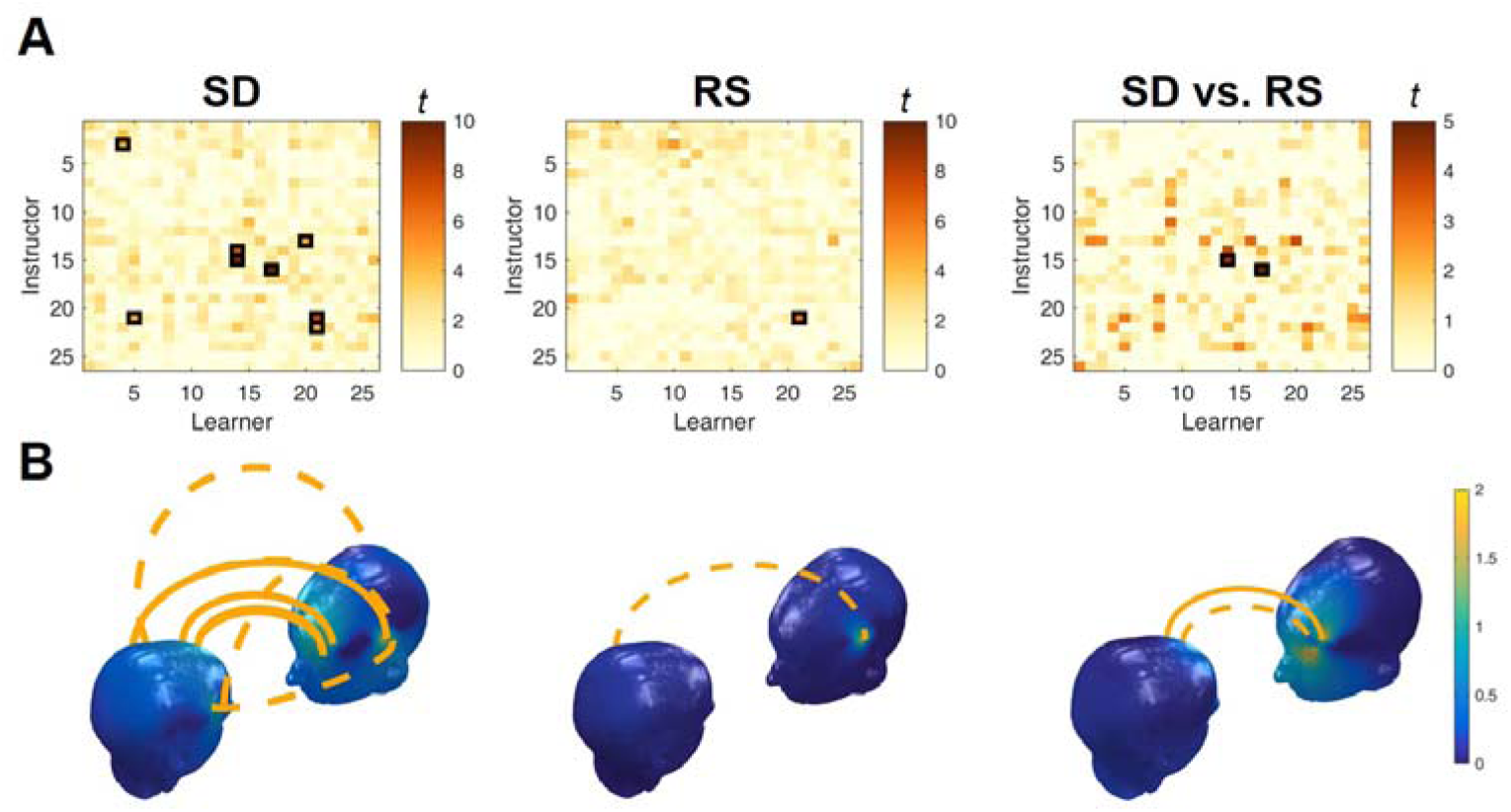
Sleep deprivation-related interpersonal brain synchronization (IBS). (A) Dissociated IBS patterns in the SD and RS conditions. SD compared to RS elicited significantly stronger IBS. The x-axis represents channels from learners whereas the y-axis represents those from instructors. The black rectangles highlight significant results thresholded at *p* < 0.05 (FDR corrected). (B) Condition-specific IBS in widespread fronto-temporo-parietal regions in the SD condition (left column) and in the superior temporal cortices in the RS condition (middle column). There was significantly higher IBS at the inferior frontal cortices in the SD than the RS condition (right column). Dash line indicates FDR corrected *p* < 0.05; solid line represents FDR corrected *p* < 0.01. Head color reflects the number of significant IBS links.

Additional analyses evidenced stronger IBS in the SD than the RS condition at channel combination CH15-CH14 (SD vs. RS, mean ± standard deviation, 0.12 ± 0.03 vs. 0.03 ± 0.08, *t*_15_ = 4.52, corrected *p* = 0.04), and CH16-CH17 (SD vs. RS, 0.13 ± 0.05 vs. 0.01 ± 0.07, *t*_15_ = 6.40, corrected *p* = 0.002; note that the first and second channel labels separated by a dash represent the instructor and learner channels that took part in the IBS). These channels roughly correspond to left inferior frontal cortex locations [75] (**Fig. 4**, right column).

Altogether, our results suggest that IBS increased in a compensatory manner during learning after SD. The effect was specific to the real interacting dyad, as computations on pseudo-randomly re-paired dyads were all non-significant.

### 3.4. Associations between (clustered) IBS and learning improvement

We next examined the relation between IBS and improvement in learning performance. There were no significant univariate correlations between the increase of IBS in either SD or RS conditions and learning improvement. To further extract meaningful information from multi-dimensional arrays (**Fig. 4**), IBS illustrating the unique instructor-learner brain-to-brain network were clustered separately in the SD and RS conditions using nonnegative matrix factorization. The three-cluster solution was used to find the best fit in the SD and RS condition (**Fig. 5A**). Cluster 2 exhibited a significant correlation with learning improvement (learning performance at the post-minus pre-learning phase) in the SD condition, *r* = 0.62, *p* = 0.01. That is, increased IBS in cluster 2 was associated with improvement in numerical reasoning performance in the learner (**Fig. 5B**). Note that cluster 2 in the SD condition mostly engaged IFC regions (i.e., CH14_CH14, CH15_CH14; **Fig. 5A**), which echoed with our above findings. Correlations with cluster 1 (*r* = -0.07, *p* = 0.80) and cluster 3 (*r* = 0.38, *p* = 0.17) were not significant. In the RS condition, no cluster was correlated with improvement in numerical reasoning performance (*r*s < 0.24, *p*s > 0.37).

**Fig. 5.**
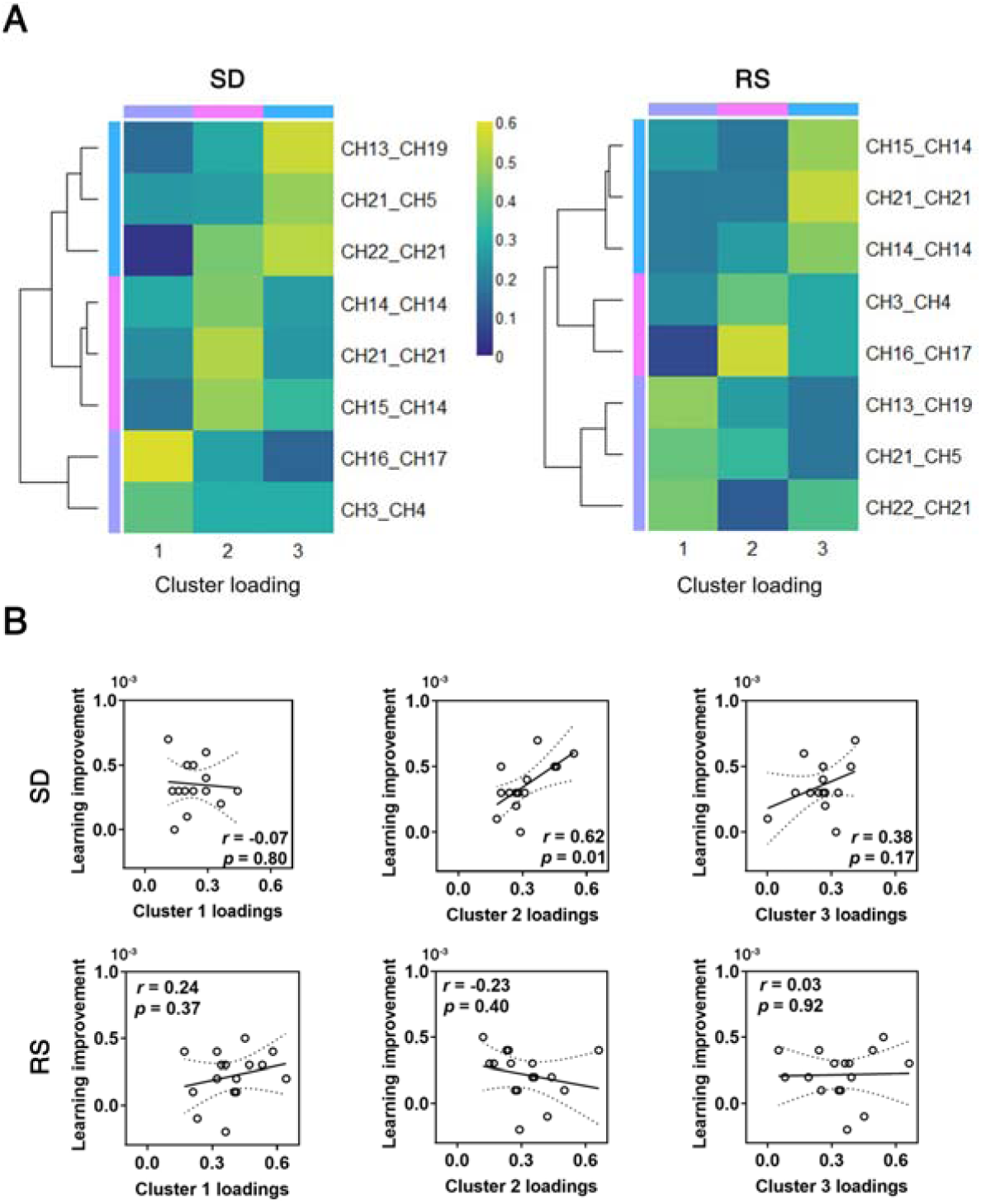
Clustered interpersonal brain synchronization (IBS). (A) Heatmaps of the three-cluster solution for significant IBS in the SD and RS conditions. The colours reflect IBS loadings for each cluster. The tree diagram illustrates the arrangement of the clusters produced by hierarchical clustering. The first and second channel names separated by an underscore respectively represent the instructor and learner NIRS channels that were involved in the IBS. (B) Scatter plots of clustered interpersonal brain synchronization (IBS) vs. learning improvement in sleep-deprived (SD) and regular-sleep (RS) conditions. Note that only Cluster 2 loadings were significantly correlated with learning improvement (calculated by post- minus pre- numerical learning performance).

We additionally probed a potential relationship between IBS changes and empathic deficit in the SD condition. Specifically, Pearson correlational analyses were conducted to test the correlations between clustered IBS and subjective/objective empathic deficits measures (delta = SD values at 9:00 AM minus RS values at 9:00 AM). Neither subjective (*r*s < 0.42, *p*s > 0.11) nor objective (*r*s < 0.31, *p*s > 0.24) measurements correlated with clusters 1–3.

### 3.5. IBS directionality: from instructor to learner or from learner to instructor?

To determine the preferential directionality of IBS during learning interactions between instructor and learner, we conducted a Granger causality analysis (GCA). Due to the data being not normally distributed, we performed a series of planned contrasts using nonparametric Wilcoxon tests. In the SD condition, Wilcoxon tests revealed a significantly biased directionality with mean causality from instructor to learner (median ± median absolute deviation, 0.0028 ± 0.0013) significantly larger than that from learner to instructor (0.0021 ± 0.0009), corrected *p* < 0.05 (**Fig. 6**). In the RS condition, there was no significant difference in terms of the coupling directionality (from instructor to learner vs. from learner to instructor, 0.0025 ± 0.0009 vs. 0.0022 ± 0.0011, *p* = 0.13; **Fig. 6**). These results suggest that it is mostly the instructor who “led” the interaction with the learner in the SD condition.

**Fig. 6.**
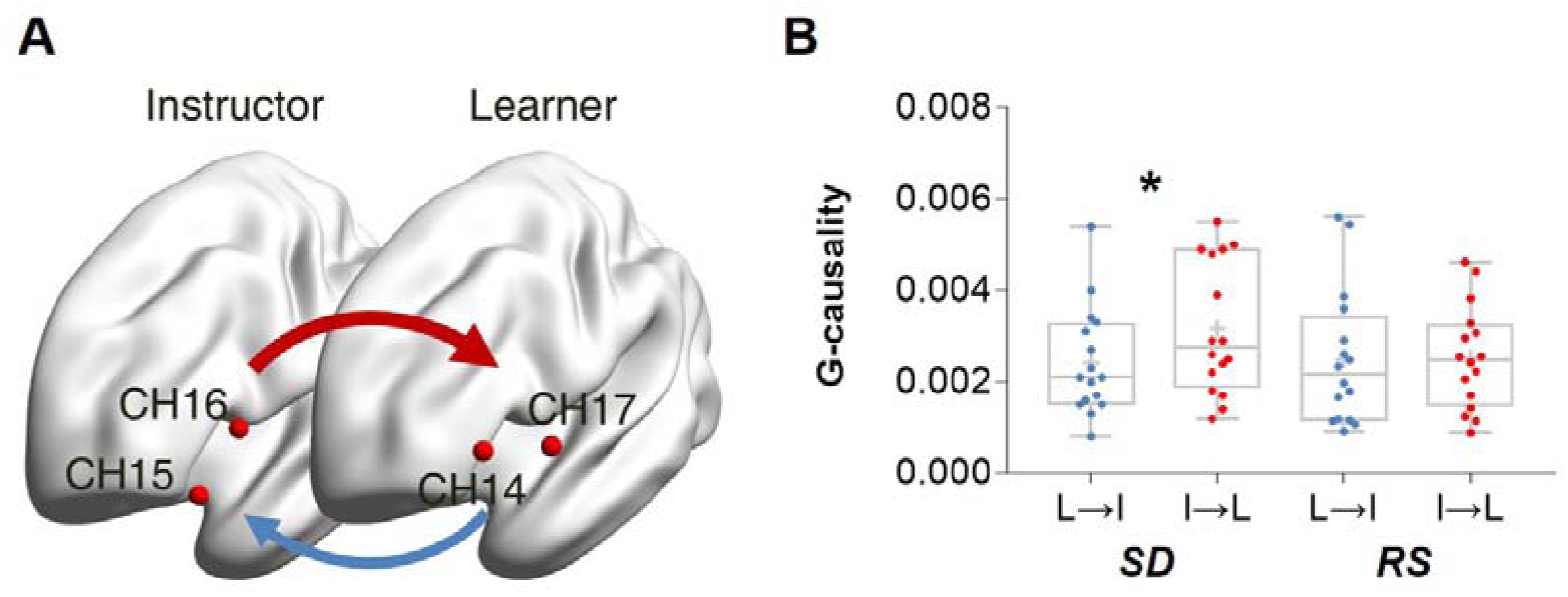
Coupling directionality. (A) ROI-based Granger causality analysis evidencing the main coupling directionality. In the SD condition, mean Granger causality from instructor [averaged signals recorded at channel (CH) 15 and CH16] to learner [averaged signals recorded CH14 and CH17] was larger than Granger causality from learner to instructor in the inferior frontal cortex. (B) In the SD condition, mean causality from instructor (I) to learner (L) was significantly larger than vice versa. In the RS condition, no significant differences were evidenced. * *p* < 0.05.

### 3.6. Synchronized seed-based intrinsic brain network after SD

As a complementary analysis, we also investigated within-individual brain connectivity during learning interactions, separately in the learner and in the instructor. In the SD condition, a seed-based intrinsic brain network analysis (**Fig. 7**) revealed significant long-range connectivity between the left inferior frontal cortex (lIFC, CH15) and the right premotor cortex (rPMC, CH6) in the instructors, *t*_15_ = 4.08, corrected *p* = 0.03, and between lIFC and right superior temporal cortex (rSTC, CH9) in the learners, *t*_15_ = 6.40, corrected *p* = 0.0003. No significant connectivity was found in the RS condition neither in the learner or the instructor. Accordingly, a between-conditions comparison (SD vs. RS) revealed significantly larger lIFC-rPMC connectivity in the SD (mean ± standard deviation, 0.07 ± 0.02) than the RS (−0.02 ± 0.02) condition for the instructor, *t*_15_ = 3.88, corrected *p* = 0.01, and significantly stronger lIFC-rSTC connectivity in the SD (0.07 ± 0.01) than RS (0.01 ± 0.02) condition for the learner, *t*_15_ = 3.98, corrected *p* = 0.01. In a third contrast (instructor vs. learner), we found that in the SD condition, interactive learning engaged larger lIFC-rPMC connectivity in the instructor (0.07 ± 0.02) than in the learner (−0.01 ± 0.01), *t*_15_ = 3.56, corrected *p* = 0.03. Considering the inter-dependencies of some data points (i.e., two instructors participated 8 times each), a multilevel mixed-effects modelling approach (the fixed effect was condition; random effects were estimated for learner and instructor, while the former was nested within the latter) was also attempted and provided analogous results. These results thus suggest that during interactive learning after SD, compensatory intrinsic synchronization took place in action-observation circuits (IFC, PMC, and STC); echoing the GCA findings that the (not sleep-deprived) instructor might play a more important role than the sleep-deprived learner in establishing the brain interactions eventually leading to successful learning.

**Fig. 7.**
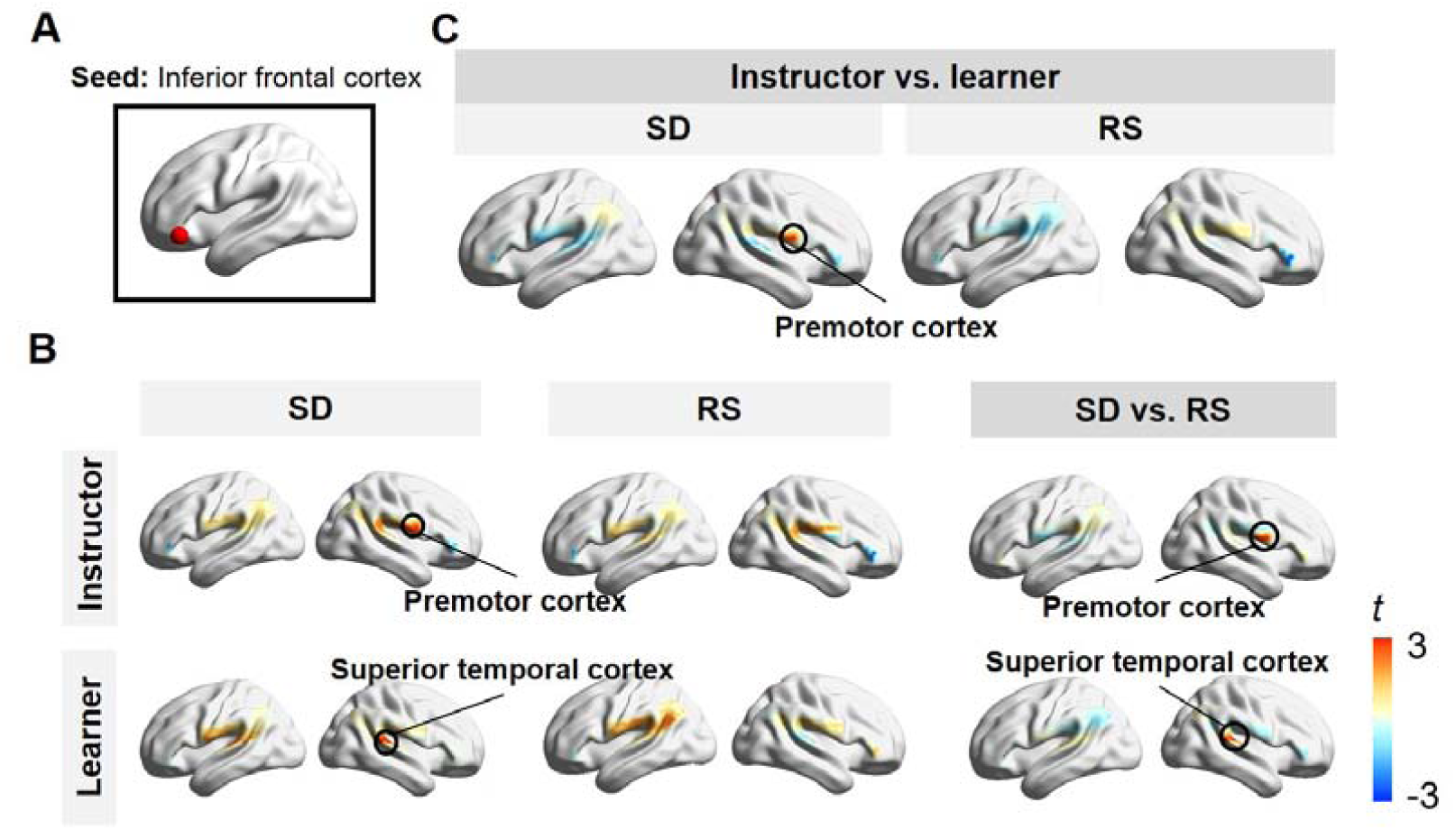
Seed-based intrinsic brain network. (A) Inferior frontal cortex as the seed region. (B) Brain regions showing task-related (task vs. rest) connectivity in the SD (left column) and RS conditions (middle column); brain regions showing higher connectivity in the SD than RS conditions (right column). (C) Brain regions showing higher connectivity for the instructor than the learner in the SD and RS conditions. Circles point to regions where connectivity was significant after FDR correction (thresholded at *p* < 0.05). Colour bars denote the *t*-value range.

## 4. Discussion

In this study, we recorded brain activity simultaneously both in instructor and learner during the interactive learning of numerical reasoning abilities. At the behavioural level, numerical reasoning performance was impaired in the learner following one night of sleep deprivation (SD). Notwithstanding, sleep-deprived learners proportionally improved to the same extent after the learning interactive session than after one night of regular sleep. At the neurophysiological level, SD compensation was characterized by increased IBS in the inferior frontal cortex (IFC) between the learner and the instructor, which was predictive of learning achievements. Further analyses revealed that enhanced IBS was mostly attributable to the instructor, as compared to the learner. Finally, increased IBS after learning was accompanied by better seed-based synchronization in intrinsic brain networks both in the learner and the instructor. These results suggest that the social interactive learning in the context of SD features enhanced IBS in the IFC, as a compensatory recruitment process.

### 4.1. Brain-to-brain coupling as a general compensatory mechanism counteracting sleep loss?

Recent studies using interactive learning tasks have shown that the instructor-learner interactions can be tracked by their IBS [26,29]. In the current study, we replicate previous findings showing that social interactions in a naturalistic environment induce synchronous activities in theory-of-mind-related brain areas including inferior frontal, superior temporal, and temporal parietal regions. Besides validating prior findings, we show that IBS and interactive learning are modulated by physiological constraints such as sleep deprivation. We surmise here that the IBS might act as a “tie”, through which the instructor continuously entrains the learner to align her/his behaviours as well as underlying activity in neural networks, facilitating social interaction and information exchange. To successfully achieve interpersonal alignment, both instructors and learners should recruit compensatory neural resources, beyond those utilized after a regular night of sleep to keep “in sync”. In this context, additional enhanced IBS eventually supports the instructor-learner interaction to successfully improve the learners’ performance. This hypothesis is supported by results from our comparison between IBS in the SD and RS conditions; as SD elicited more widespread IBS than RS. Moreover, we found a significant association between clustered IBS (represented by cluster 2) and learning improvement after SD, suggesting functional significance in the IBS. Besides individual SD-related compensatory recruitment [17,20], our results suggest that compensation after SD could impact both interacting partners simultaneously, even if one of them was actually not sleep-deprived. Noticeably, IBS was not significantly associated with learning outcomes in the RS condition, at variance with previous studies that motivated our paradigm [26,29]. This lack of relationship deserves further exploration, as it may reflect either a lack of sensitivity of continuous IBS analysis [28] or a reduced interaction between learners and instructors in a less challenging well rested condition.

One may argue that the unraveled IBS simply reflect the functional similarities between two brains processing the same sensory information or performing the same actions simultaneously [37,76]. We argue that this is not the case in the present study for the following reasons. First, SD induced larger and more widespread IBS compared to RS, although both conditions shared similar instructions and sensory inputs. Second, the pseudo-dyad control analysis further excluded the potential confound of non-interaction behaviors on the increase of IBS (i.e., it was not expected that IBS emerged in pseudo-dyads since they performed similar task but produced non-interaction behaviors). Finally, strongly highlighting the functional significance of IBS, we observed that the SD-related clustered IBS was positively correlated with the learning performance of these learners. Taken together, these results make it less likely that IBS emerged as a consequence of the similarity between sensory processes across the participants forming each dyad.

### 4.2. IFC as a neural communication interface between instructors and learners

In the present study, we found IBS mostly present over the inferior frontal cortex (IFC). IBS in the IFC was reported in fNIRS-based hyperscanning studies involving face-to-face communication [77], cooperative singing [78], and instructor-learner interaction situations [26]. Our results are also in agreement with available evidence that compensatory recruitment requires partially successfully behavioural adaptation, partly subtended by the IFC [17]. We here highlight three possible functional meanings of the IFC. First, the IFC is viewed as an important hub of the mirror neurons system [79], proposed to promote social interaction by predicting other individuals’ actions and intentions [80]. Thus, interactive learning could have been facilitated by the mutual abilities to infer and understand each other’s behaviour (i.e., high-level mentalizing [81]). Second, the IFC is also known as a critical language hub of the human brain. In this respect, left IFC dominance in this study might be considered alongside the syntax information parsing in the current numerical reasoning task [82]. Although linguistic and mathematical syntax in the human was reported to be independent [83], the syntax of mathematics may be evolutionarily derived from that of language or vice versa [84]. Therefore, it is possible that numerical reasoning and language share neural representations in the left IFC. Finally, related to the previous point, the effect in the left IFC could also be explained by the simpler linguistic exchanges (i.e., oral communication) between participants [77]. Clearly, more work is required to investigate the exact functional significance of the IFC during interactive learning.

Synchronized IFC (seed)-based intrinsic brain network activity paralleled IBS findings. Specifically, SD induced a long-range increase of intra-brain connectivity in the action-observation network (involving inferior frontal, premotor, and superior temporal cortices), which was proposed to support the understanding of other individuals’ goals and actions [85]. In this network, the premotor cortex and the superior temporal cortex would support motor planning and action intentionality encoding, respectively [85].

### 4.3. Instructor-entrainment as an essential feature of interactive learning following SD

A major peculiarity of our design is that we investigated synchronous brain activity between a learner, who was sleep-deprived, and an instructor, who did not receive sleep disruption at all. Granger causality analyses showed that after SD, mean causality was significantly larger from the instructor to the learner than the other way around. This suggests that the instructor might play a more important role in social interactive learning than the SD learner. As such, dynamic social interactions [86] that are key components for grasping the others’ mind would play an important role in sleep-deprived interactive learning. Following SD, despite brain activity compensation in the learner, the instructor guided the communication, monitored the learner’s responses, and entrained her/his brain activity with the one of the learners. This interpretation is reinforced by a seed-based intrinsic brain network analysis showing that the instructors displayed additionally and significantly stronger intra IFC-PMC connectivity than the learners. It suggests that the instructor actively guide the learner and help her/him to counteract the effects of sleep disruption. Taken together, instructor-entrainment might be a marker of the essential nature of interactive learning following SD, as it is much more prominent than when the learner is in a rested state.

As a coin has two sides, an increase in how the instructor drives the learner’s brain activity could also be interpreted as an increase in the learner’s passivity. In this respect, one may argue that the increase in IBS or directionality is not a necessarily positive outcome. However, we found additional IFC-STC intrinsic brain connectivity in learners in the SD condition compared to the RS condition. This result partially opposes the “learner’s passivity” hypothesis in interactive learning following SD.

### 4.4. Limitations

This study has several limitations. First, we investigated only the effects in female dyads in order to mitigate inter-individual and inter-dyad variability [26,37]. It remains unclear whether the findings could be generalizable to males. There could be gender difference given that males and females may differ in dyadic interaction [87]. Second, we had the learners sleep at home rather than in the laboratory the night before testing in the RS condition. This practice was to respect naturalistic conditions at maximum while controlling sleep schedules by the instruction and actigraphy (see Methods). Yet, the SD and RS conditions differed in the time spent to get familiar with the testing environment. Third, the effect of SD might be confounded by cognitive fatigue. For example, participants took sleepiness and empathy assessments multiple times in the SD condition but took these assessments only once in the RS condition. Recent advances showed that cognitive fatigue and sleepiness can be dissociated when accumulated sleep pressure is low [88]. Future studies should further examine and control the confounding effect from cognitive fatigue in SD interactive learning.

### 4.5. Conclusions and future directions

In the current study, we have evidenced the benefits of combining an interactive learning paradigm in educational psychology with an fNIRS hyperscanning paradigm used in social neuroscience. Whereas interactive learning allowed us to study the educational interaction between an instructor and a learner in a realistic context, the fNIRS hyperscanning approach enabled quantifying inter-individual information flows and directionality at the neurophysiological level. Although we manipulated only a single interaction situation (i.e., interactive learning), our study may promote future studies to investigate related learning and educational issues. In addition, our findings of compensatory IBS and intrinsic network connectivity primarily are in line with the hypothesis of compensatory brain recruitment to at least partially counteract the deleterious effects of SD in a learner. Future studies could consolidate the current findings by adding a “single” condition as an active control, in which instructor and learner would independently solve numerical reasoning problems. In this case, they would be both intentionally thinking about the same thing but without the interpersonal interaction component.

To sum up, our study shed light on whether and how learning performance could be compensated in interactive learning after that a learner experienced one night of sleep deprivation. Our results suggest that brain-to-brain coupling, accompanied by synchronization in intrinsic brain networks, may require the involvement and integration of high-level social cognitive processes to compensate for interactive learning following SD, and that the instructor might play a vital role in driving synchrony between teaching and SD learning brains.

## Acknowledgments

The authors would like to thank Dr. Veronica Guadagni and her colleagues for sharing the materials of the Multifaceted Empathy Test, Dr. Pavel Goldstein for helping with the nonnegative matrix factorization analysis, Dr. Guillaume Dumas for sharing the function used to visualize the brain-to-brain coupling. This work was supported by a Belgian National Fund for Scientific Research (FRS-FNRS) grant T.109.13, the National Natural Science Foundation of China (31872783), the China Scholarship Council (201706140082), and the Outstanding Doctoral Dissertation Cultivation Plan of Action of East China Normal University (YB2016011).

